# Haptoglobin buffers lipopolysaccharides to safeguard against aberrant NFκB activation

**DOI:** 10.1101/2022.11.22.516855

**Authors:** Laura Zein, Josina Grossmann, Helena Swoboda, Christina Borgel, Bernhard Wilke, Stephan Awe, Andrea Nist, Thorsten Stiewe, Oliver Stehling, Sven-Andreas Freibert, Till Adhikary, Ho-Ryun Chung

## Abstract

It has remained yet unclear which soluble factors regulate the anti-inflammatory macrophage phenotype observed in both homeostasis and tumourigenesis. We show here that haptoglobin, a major serum protein with elusive immunoregulatory properties, binds and buffers serum-borne bacterial lipopolysaccharides to attenuate activation of NFκB in macrophages. Haptoglobin binds different lipopolysaccharides with low micromolar affinities. Given its abundance, haptoglobin constitutes a buffer for lipopolysaccharides, shielding them to safeguard against aberrant inflammatory reactions caused by small fluctuations. Concordantly, NFκB activation by haptoglobin-associated lipopolysaccharides was delayed relative to stimulation with pure lipopolysaccharide. Our findings warrant evaluation of therapeutic benefits of haptoglobin for inflammatory conditions and re-evaluation of purification strategies. Finally, they allow to elucidate mechanisms of enhanced immunosuppression by oncofetal haptoglobin.

## Introduction

Macrophages are cells of the innate immune system which control inflammation, wound healing, and homeostasis. The abundance of tumour-associated macrophages (TAMs) is correlated with poor prognosis in several tumour types [1]. We compared *ex vivo* (freshly isolated) TAMs from ascites of ovarian carcinoma (OC) with *ex vivo* peritoneal macrophages from tumour-free patients and found them to be highly similar; both display a predominantly immunosuppressive phenotype according to RNA-seq and flow cytometry. Discernible differences are limited to (i) TAM quantity—vastly outnumbering other immunocompetent populations—and (ii) expression of rather small sets of genes. These sets contain either pro-tumourigenic genes involved in extracellular matrix remodeling, which is a hallmark of the resolution phase of inflammation that is also observed in wound healing, or genes regulated by anti-tumourigenic interferon signalling. In contrast, we found that macrophages differentiated *in vitro* have very different transcriptomes [2, 3], showing that *in vitro* polarisation does not recapitulate the *ex vivo* state. It is therefore of particular interest which factors are present *in vivo* that modulate expression of inflammation-related genes to shift the balance between pro- and anti-inflammatory macrophage phenotypes in health and disease.

The NFκB pathway regulates immune cell function. NFκB signalling is stimulated by pathogen recognition receptors (PRRs), such as the Toll-like receptors (TLRs), and by other receptor families including specific cytokine receptors. TLR4 is the archetypical PRR. TLR4 is expressed on monocytes, macrophages, dendritic cells, B cells, adipocytes, endothelial cells, and on Paneth cells of the intestinal epithelium. Together with its co-receptors CD14 and MD2, TLR4 activates the NFκB pathway after binding of its agonist, lipopolysaccha-ride (LPS). NFκB signalling culminates in phosphorylation of inhibitor of κB (IκB) proteins, their subsequent ubiquitination and degradation. This frees transcriptional activators referred to as NFκB which then translocate to the nucleus and induce transcription of their target genes. The temporal dynamics of NFκB nuclear translocation encode for ligand and dose to determine biological responses [4]. NFκB targets include many pro-inflammatory genes, but NFκB also regulates development and homeostasis [5]. A major subset of NFκB target genes in macrophages, including *IL1B, IL12B*, and *TNF*, is involved in pro-inflammatory processes, whereas another major subset exemplified by *IL6, IL10*, and *PTGS2* (encoding for cyclooxygenase 2) mediates immunosuppression in homeostasis, wound healing, and neoplasia. Most NFκB target genes including *IL1B, IL6, IL10*, and *PTGS2* are expressed in OC TAMs and in peritoneal macrophages [2, 3, 6]. This raises the question which endogenous factors regulate NFκB signaling in macrophages *in vivo* to achieve an anti-inflammatory, homeostatic macrophage polarisation state.

Haptoglobin (HP) is an acute phase protein with concentrations of 0.3–2 g/l in adult human serum [7]. HP sequesters free hemoglobin (HB) to prevent oxidative tissue damage upon hemolysis [8]. Conflicting HB-independent functions of HP were reported. On the one hand, HP represses LPS-induced cytokine expression [9, 10], suggesting attenuation of nuclear factor NFκB. Moreover, haptoglobin-deficient mice are prone to autoimmune inflammation [11], which implicates an anti-inflammatory function of haptoglobin. On the other hand, HP was reported to activate TLR4-NFκB signalling [12, 13]. Taken together, it has remained yet unclear how HP regulates inflammatory processes qualitatively as well as mechanistically.

LPS, alternatively called endotoxin, is an outer membrane component of Gram-negative bacteria. LPS molecules are large and heterogeneous glycans composed of a lipid A moiety, a core oligosaccharide moiety, and a repeating polysaccharide O antigen. The human gut contains approximately 1 g of LPS. Intestinal permeability allows LPS to traverse into the bloodstream [14, 15], and LPS is present in amounts of 1– 200 pg/ml in human serum [14]. High-fat meals are known to induce endotoxemia and inflammatory markers [16]. Notably, elevated LPS concentrations in human serum have been reported in obesity and diabetes [17], ethanol abuse [18], and neurodegenerative disorders [19]. In animal models, exposure to LPS induces obesity, diabetes, and neurodegeneration [20]. The involvement of LPS in the genesis of cancer has been implicated frequently [21]. These data collectively suggest that endotoxemia is causal for different pathophys-iologies. Notably, LPS molecules from bacterial species and strains differ in their molecular composition, and some do not activate NFκB. This is exemplified by LPS-Rs from *Rhodobacter sphaeroides*, which antagonises TLR4-dependent NFκB activation by other LPS species. The molecular basis for these observations is that TLR4 binds to the lipid A moiety invariably present in all LPS molecules which hence is the immunogenic part of LPS.

*Per se*, LPS is not a toxin; it elicits a TLR4-mediated cytotoxic host response in mammals. LPS availability needs to be tightly controlled to prevent acute inflammation. Proteins which specifically bind to LPS include soluble CD14, LPS-binding protein (LBP), BPI, APOE, adiponectin, α-defensins, surfactant proteins, and lactoferrin [22]. Genetic deletion of LBP causes susceptibility to endotoxemia in mice [23]. The need for adequate buffering of LPS was proposed [22]. The presence of endogenous stores, dedicated carriers, the TLR4 receptor complex, and mechanisms which specifically counteract the response to LPS in mammals underscores the involvement of LPS in homeostasis and resulted in designation of LPS as an exogenous hormone [24]. This is in line with the unique role of TLR4: It is the only TLR that recruits all four adaptor proteins MYD88, TIRAP, TRAM, and TRIF to elicit a distinct gene expression profile [25].

Here, we show that the conflicting functions of HP in TLR4-NFκB signalling are explainable by HP’s ability to bind and buffer LPSs, which results in shielding of LPS from TLR4 and delayed NFκB activation.

## Results

HP purified from human serum induced the expression of the NFκB target gene *IL1B* (>1,000-fold; fig. 1A) in monocyte-derived macrophages (MDMs). By contrast, HB alone did not induce *IL1B* expression (fig. 1A), which together with the fact that the HP preparation contained only spurious amounts of HB (fig. 1B) indicates that HB is dispensable for *IL1B* induction by HP. Transcriptome analysis of HP-treated *versus* control MDMs identified differentially expressed genes that are typical for an NFκB-dependent response (fig. 1C).

**Figure 1:**
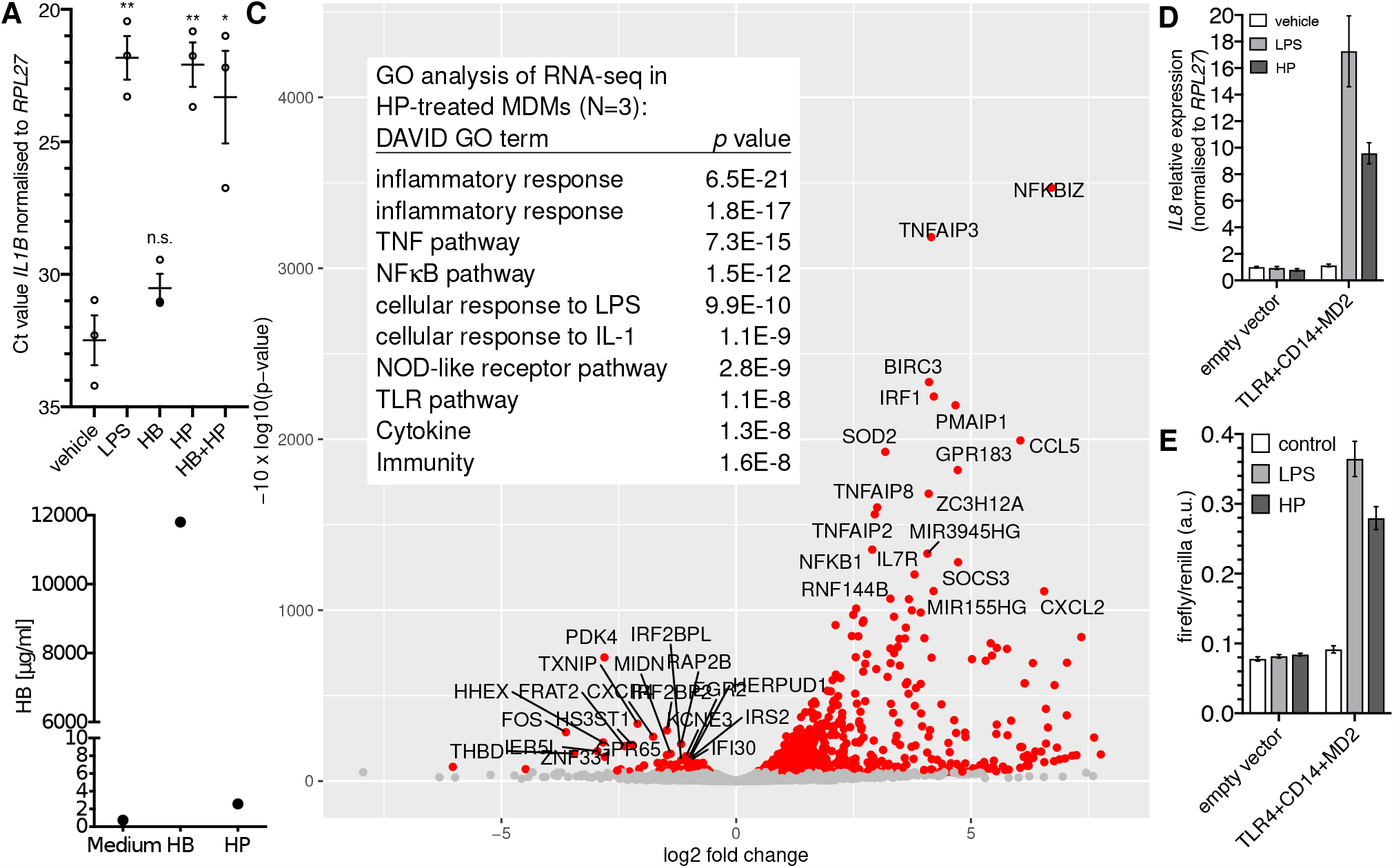
Purified haptoglobin induces NFκB-dependent transcription through TLR4. (**A**) MDMs from blood donors (N=3) were treated with 100 ng/ml *E*.*coli* LPS or HB (25 μg/ml), mixed type HP (25 μg/ml), or both for six hours. Expression of *IL1B* was measured by RT-qPCR. Error bars represent standard deviations. Bonferroni-corrected significance (unpaired *t* test): ^**^, p<0.01; ^*^, p<0.05; n.s., not significant. (**B**) ELISA analyses of the hemoglobin content of HB and HP. Nominal protein concentration is 10 mg/ml each. (**C**) Volcano plot of RNA-seq data; HP treatment of MDMs from three donors (HP; 25 μg/ml for 4 h *vs*. solvent control). Table inlay: Top ten GO terms assigned by the DAVID database to the top 50 upregulated genes. (**D**) HEK293 cells were transfected as indicated and treated with either solvent, *E*.*coli* LPS (100 ng/ml), or HP (25 μg/ml) for 6 h. *IL8* expression was monitored by RT-qPCR. This is representative of three independent experiments. (**E**) HEK293 cells were transfected as in A plus an NFκB firefly luciferase reporter plasmid (5×NFκB-luc) and a constitutive *Renilla*-luc reporter and treated as in A. Representative of two independent experiments. D, E: Error bars represent standard deviations from three technical replicates.

The NFκB-type transcriptomic response led to the idea that HP activates NFκB *via* TLR4. To test this, we performed a synthetic complementation assay in HEK293 cells, where only the forced expression of TLR4, CD14, and MD2 led to induction of *IL8* by purified HP (fig. 1D) indicating that TLR4, CD14, and MD2 together are sufficient to confer sensitivity to HP. In line with this idea, an NFκB-responsive reporter was induced by HP only when we complemented TLR4, CD14, and MD2 in HEK293 cells (fig. 1E), demonstrating that purified HP activates NFκB-dependent transcription *via* TLR4.

Since HP binds to TLR4 [13], we reasoned that HP directly activates NFκB-dependent transcription *via* TLR4. To test this hypothesis, we enzymatically digested HP using proteinase K. Unexpectedly, we found that proteolysis did not abolish NFκB-dependent transcription (fig. 2A). Thus, HP protein is dispensable for the activation of NFκB-dependent transcription *via* TLR4.

**Figure 2:**
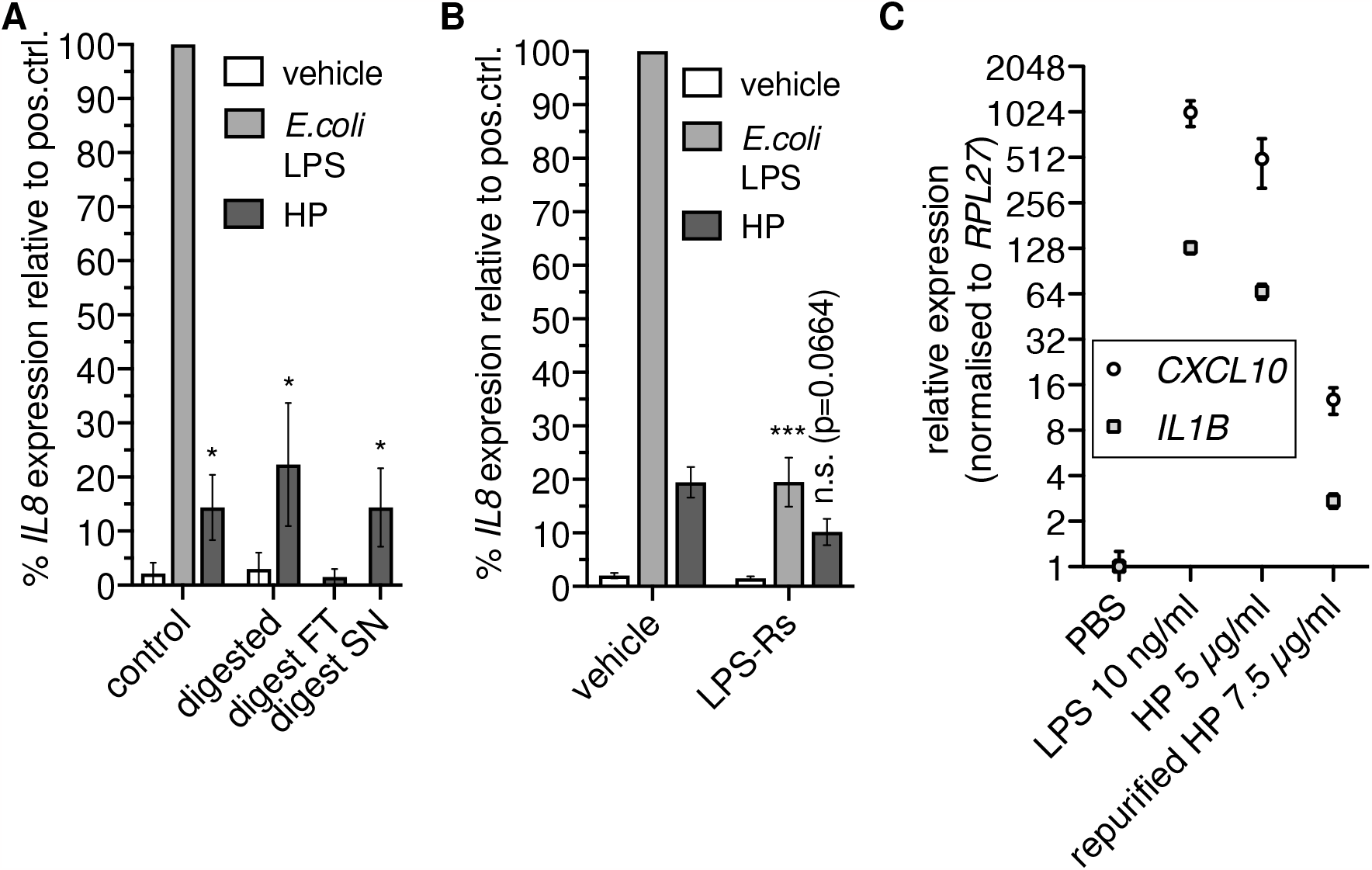
Haptoglobin is dispensable for NFκB-dependent transcription. (**A**) Mixed type HP was treated with proteinase K. An aliquot was ultrafiltrated with a 10 kDa cutoff; FT, flowthrough; SN, supernatant. THP-1 cells were treated for 6 h with these preparations as indicated or with 100 ng/ml *E*.*coli* O111:B4 LPS, and RT-qPCR of the *IL8* transcript was performed. ^*^, *p<*0.05 relative to the corresponding negative control (unpaired *t* test). Error bars represent SD. One of two independent experiments is shown. (**B**) THP-1 cells were treated as in A with or without LPS-Rs (10 μg/ml). Error bars represent SEM (N=6). ^***^, *p<*0.001; ^*^, *p<*0.05 relative to the corresponding sample without LPS-Rs (unpaired *t* test). (**C**) THP-1 cells were treated for 6 h with *E*.*coli* LPS, HP, or HP repurified by gel filtration in the presence of 500 mM NaCl, and RT-qPCR of the *CXCL10* and *IL1B* transcripts was performed. Error bars represent SD from three technical replicates.

The stimulus remained in the supernatant after ultrafiltration of protease-treated samples with a 10 kDa cutoff (fig. 2A). We then checked whether the TLR4 antagonist LPS-Rs (from *R*.*sphaeroides*) competes with the non-protein factor in the HP preparation. LPS-Rs led to reduced *IL8* induction in response to HP (fig. 2B), indicating the presence of TLR4 agonists. To further substantiate the claim that co-purified TLR4 agonists rather than the HP protein activate NFκB-dependent transcription *via* TLR4, we treated THP-1 cells with HP protein repurified by gel filtration under high-salt conditions. Repurification abrogated induction of NFκB target genes by HP (fig. 2C). Together, these data indicate that not HP itself but a separable, non-protein TLR4 agonist is responsible for induction of NFκB target gene expression by HP preparations.

The canonical agonists of TLR4 are LPSs; we therefore speculated that HP is associated with LPSs. To test this, we used a sensitive silver stain to detect LPSs [26] in digested HP. The assay revealed distinct high molecular weight bands, which are regularly observed in LPS preparations from different bacteria (especially in clinical isolates [27]) and indicate the presence of long-chain “smooth” LPSs (fig. 3A). Slower migration of discernible leading bands suggests that the lipid A and inner core moieties of the LPSs differ from the *E*.*coli* reference, which exhibits a ladder of regularly increasing chain lengths. Thus, the detected LPSs largely originate from various bacterial species other than *E*.*coli*.

**Figure 3:**
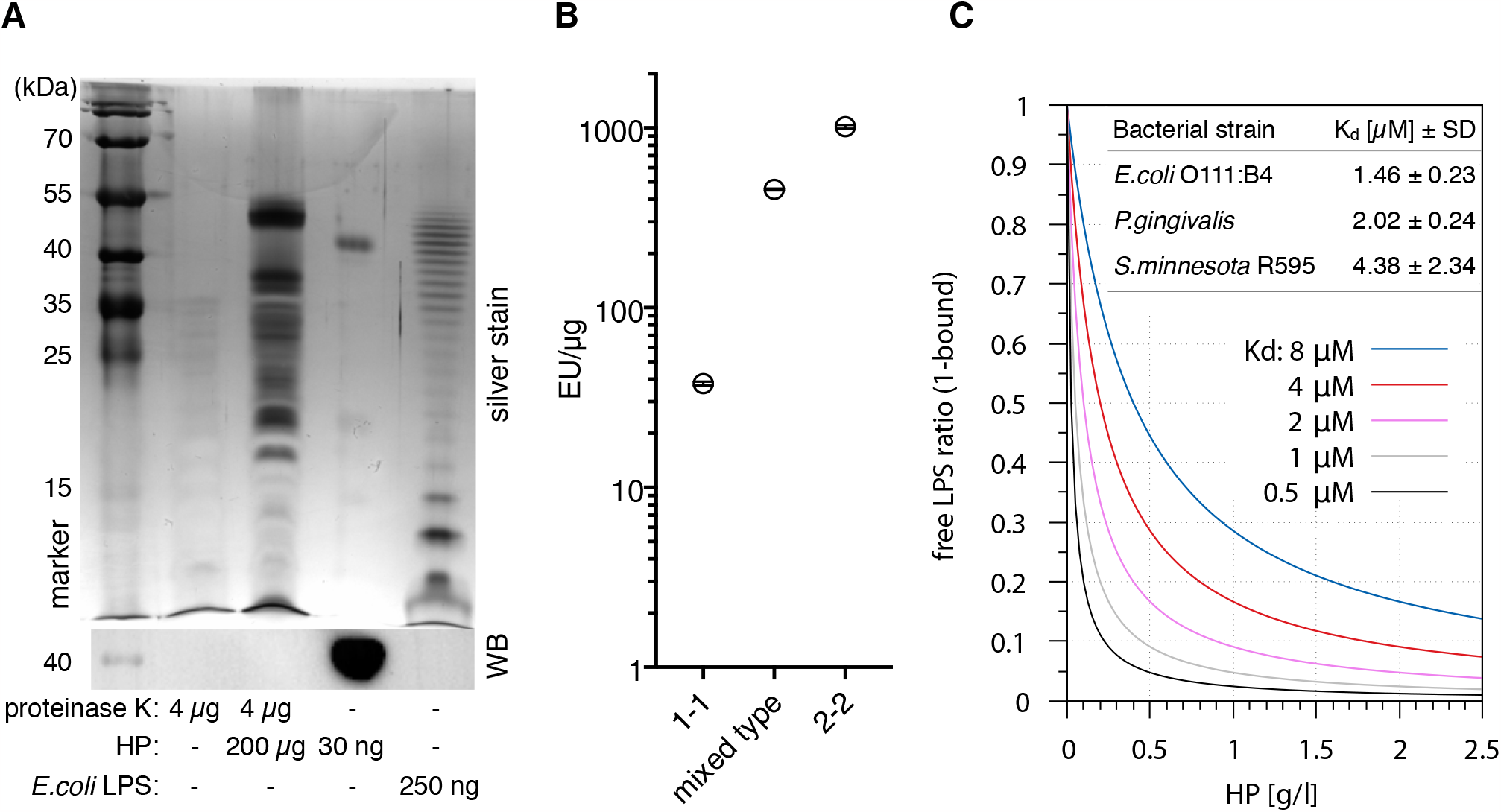
Haptoglobin is associated with and binds lipopolysaccharides. (**A**) Upper panel: Silver stain of the indicated samples after deoxycholate-urea PAGE under reducing conditions. HP was digested with proteinase K for 2 h at 55 °C or not as indicated. Digestion of the protein was complete as indicated by immunoblotting against the HP β chain (lower panel). (**B**) A LAL assay was conducted with HP1-1 (from homozygous carriers of allele 1, resulting in dimer formation), HP2-2 (oligomeric HP from homozygous carriers of allele 2), and mixed type HP from heterozygous carriers (N=1 each). Error bars represent SD calculated from three technical replicates. EU, endotoxin units. (**C**) Calculated ratios of free *vs*. HP-bound LPS at the indicated *K* d values. Physiological HP concentrations range from 0.3–2 g/l. At 1 g/l, HP concentration per αβ-subunit is 20 μM. Table inlay: MST data obtained with mixed type HP repurified using gel filtration and ultrapure LPS preparations from the indicated bacterial strains. MST measurements were performed with three HP preparations each.

Two main *HP* alleles exist in humans. HP from allele 1 harbours one multimerisation domain, and homozygous carriers express HP1-1 dimers. Allele 2 encodes for two multimerisation domains due to a duplication of two exons, resulting in oligomer formation [7]. A *Limulus* amoebocyte lysate (LAL) assay detected 40–1000 endotoxin units (EU)/μg protein in HP preparations of the three isotypes (dimeric HP1-1, oligomeric HP2-2, and mixed type). One EU is equivalent to 0.1–0.2 ng LPS—at 1 mg/ml HP in serum, this translates into >4 μg/ml LPS, which is >1.000-fold above reported serum LPS levels of 1–200 pg/ml [14] but in line with the strong staining we observed (fig. 3B). Since the LPSs are heterogeneous and different from *E*.*coli* LPS (fig. 3A), contamination of the HP preparations seems unlikely. A more parsimonious explanation for large amount of LPSs in the preparations is given by the fact that LPS and LPS-bound proteins are selectively precipitated by ethanol [28, 29]. It is conceivable that the widely used Cohn cold ethanol serum fractionation protocol leads to enrichment of LPS-bound HP.

Microscale thermophoresis (MST) with LPS-free HP (repurified by gel filtration in high-salt buffer) indicates *K* _d_ values <10 μM for LPSs from three bacterial species (fig. 3C) that represent different lipid A structures [30]. The *S*.*minnesota* strain R595 produces rough LPS (lacking the repeating oligosaccharide units O antigen) exclusively; binding of this molecule establishes that HP interacts with the lipid A or inner core moiety (or both). The law of mass action dictates that the bulk of LPS in serum is bound by HP in the physiological range of HP levels—1 g/l of HP corresponds to a molarity of 20 μM per HP αβ subunit, which is well above the *K* _d_ values we obtained for different LPSs. The fraction of HP-bound LPS at different values of *K* _d_ is illustrated in fig. 3C.

These results show that commercially available HP contains LPSs such that the addition of these HP preparations to cells leads to the release of LPSs, which in turn activate TLR4. However, activation should be delayed due to limited LPS availability: HP competes with TLR4 for LPS. In line with this notion, we found that IκBα degradation induced by HP was delayed relative to high-dose and low-dose *E*.*coli* LPS (fig. 4). IκBα degradation kinetics depend on LPS concentration [31, 32]. Although the deployed HP should contain an amount of LPSs in the range of the *E*.*coli* LPS controls, IκBα degradation was much slower, indicating that only a comparably low amount of LPSs was released.

**Figure 4:**
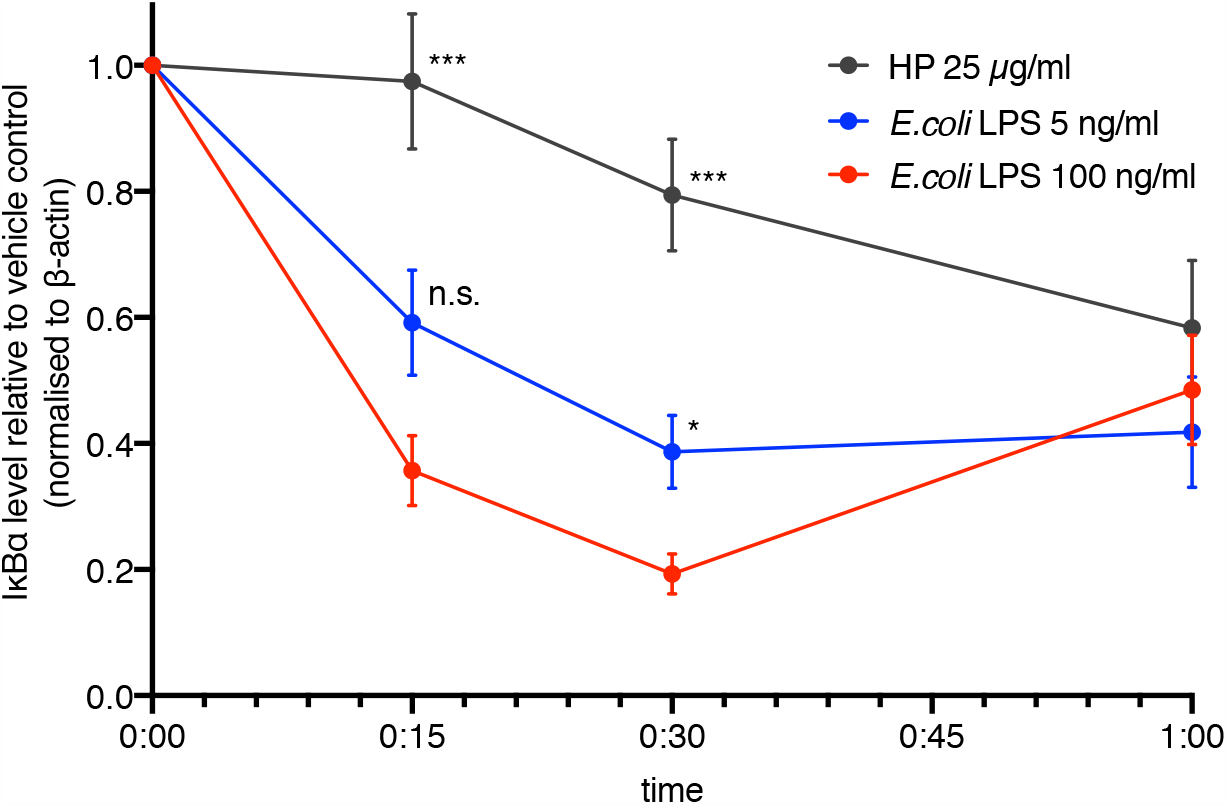
NFκB activation is delayed in the presence of haptoglobin. MDMs were treated with *E*.*coli* LPS (100 ng/ml, 5 ng/ml) or mixed type HP (25 μg/ml). IκBα levels were monitored by immunoblotting. Chemoluminescence was quantitated with a CCD-based imaging system. IκBα levels were normalised to β-actin. Dots indicate the calculated means (N=7). Error bars denote SEM. Significance relative to high-dose LPS (unpaired *t* test): ^***^, *p<*0.001; ^*^, *p<*0.05; n.s., not significant. Significance was Bonferroni-corrected for multiple hypothesis testing.

## Discussion

We show that HP purified from human serum binds LPSs with low micromolar affinities. Our data thus provide a mechanistic explanation for conflicting observations on the role of HP in NFκB signaling [9, 10, 12, 13]. Moreover, they establish HP to function as a buffer for LPSs. This buffering function is relevant because the rate of change of stimulus concentration controls the NFκB response [4, 32, 33]. HP dampens variations in LPS concentration by shielding LPS from TLR4, such that larger concentration changes are required to trigger an equivalent response. In agreement with this model of competition for LPS between HP and TLR4, HP reduces LPS-dependent pro-inflammatory cytokine expression in a dose-dependent manner [9].

HP is used therapeutically for hemolytic conditions [34]. Our findings extend applications to inflammatory states induced by elevated LPS levels in chronic conditions such as neurodegeneration [20], psychiatric diseases, inflammatory bowel disease, and metabolic syndrome [15] as well as acute inflammation. It is note-worthy that the SARS-CoV2 spike protein binds to LPS and enhances the TLR4-dependent inflammatory response [35, 36], and poor outcome in COVID19 is connected to elevated LPS levels [37, 38]. Increasing LPS buffering capacity by HP may therefore improve clinical outcomes in chronic and acute inflammation, COVID19, and other infectious diseases to limit deregulated LPS-dependent cytokine expression. Importantly, HP isolation procedures should avoid LPS enrichment.

The HP precursor expressed from allele 2, zonulin, increases intestinal permeability [39]. Our findings raise the possibility that zonulin binds LPS. HB binds LPS and exacerbates its effects [29, 40]. Attenuation of LPS-dependent effects by HB-binding proteins like hemopexin [10, 41] and HP [9, 10] may be a recurrent antagonistic theme. Likewise, hemopexin may directly bind LPS.

We found that HP purified from human serum is associated with LPS regardless of its oligomerisation state (fig. 3B) but consistently found less LPS associated with HP1-1 also in preparations purchased earlier (data not shown). This might indicate a lower affinity of dimeric *vs*. oligomeric HP towards LPS.

Elevated haptoglobin levels are negatively correlated with survival in ovarian carcinoma [42] and other solid tumours [43]. “Oncofetal” HP, which is observed during neoplasia and pregnancy [7], is a much stronger immunosuppressant than normal adult HP [44]; the mechanistic basis of enhanced immunosuppression remains unclear. Our findings suggest that the alternative glycosylation of oncofetal HP [45] potentially alters its affinity towards LPSs. Alternatively, oncofetal HP may regulate inflammation through LPS-independent mechanisms, which if true warrants separation of beneficial from malignant functions of HP.

## Materials and Methods

### Cells and reagents

Buffy coats from anonymised healthy female donors were from the University Clinic Marburg blood bank. MDMs were differentiated as described [46]. Mixed type HP (used for all experiments unless indicated otherwise) was from Sigma and USBio. HP1-1 and HP2-2 were from Sigma. Ultrapure LPSs from *E*.*coli* O111:B4, *R*.*sphaeroides*, and *S*.*minnesota* R595 were from Invivogen. Proteinase K was from Bioline. Amicon filters were from Merck-Millipore. The hTLR4 expression vector (Addgene 13086) was a gift from Ruslan Medzhitov. pcDNA3-CD14 (Addgene 13645) and pFlag-CMV1-hMD2 (Addgene 13028) were gifts from Doug Golenbock. pGL4.32[luc2P/NFκB-RE/Hygro] was from Promega. Antibodies were from Novus (α-HP JM10-79), Santa Cruz (α-IκBα sc-371), and Sigma (α–β-actin AC-15). The HB ELISA was from Bethyl (E88-134). The LAL chromogenic endotoxin assay kit was from Pierce (A39552).

### Expression analyses

RT-qPCR and immunoblots were performed as described [46]. Primer sequences are:

*CXCL10*: AAGCAGTTAGCAAGGAAAGGTC GACATATACTCCATGTAGGGAAGTGA

*IL1B*: TGAAAGCTCTCCACCTCCAGGGACA GAGGCCCAAGGCCACAGGTATTTTG

*IL8*: AGCTCTGTGTGAAGGTGCAGT GATAAATTTGGGGTGGAAAGGT

*RPL27* : AAAGCTGTCATCGTGAAGAAC GCTGTCACTTTGCGGGGGTAG

RNA was isolated using TRIfast [46] (Peqlab) with pre-isolation *D*.*melanogaster* Schneider S2 spike-in (1:10) and post-isolation ERCC spike-ins (Thermo Fisher) according to the manufacturer’s instructions. Libraries were prepared with QuantSeq FWD (Lexogen). Sequencing was performed on a NextSeq 550 (Illumina).

### Size exclusion chromatography and affinity measurements

1 mg HP was run on a Superdex 200 Increase 10/300 gel filtration column (Cytiva) with 500 mM NaCl in phosphate-buffered saline (PBS) on an Äkta Purifier 10 high-performance liquid chromatography system (GE Healthcare) to remove associated LPSs. The main protein peak fraction (at λ=280 nm) was used for covalent labeling with RED-NHS dye, and MST was performed as published [47] with freshly labeled protein in PBS supplemented with 0.005 % (v/v) Tween-20. Each LPS preparation was titrated in a 16-step 1:2 dilution series starting with a final assay concentration of 3.75 g/l. Average molecular weights for the LPS preparations were estimated after silver staining (*E*.*coli* O111:B4, 25 kDa; *P*.*gingivalis*, 30 kDa; *S*.*minnesota* R595, 2.5 kDa). A molecular weight of 50 kDa per HP αβ-subunit was assumed.

### Statistics

Unpaired, two-tailed *t* tests were used to calculate *p* values. Multiple hypothesis testing corrections were applied as indicated.

## Data availability

RNA-seq data and associated experimental and bioinformatic details are deposited at Gene Expression Omnibus (GSE215916).

## Acknowledgements

We gratefully acknowledge the contribution of the Protein Biochemistry and Spectroscopy Core Facility (Institute of Cytobiology, Center for Synthetic Microbiology, Philipps University Marburg) for help with MST analyses. We are grateful to Sovana Adhikary, Matthias Lauth, and Alexander Visekruna for critical reading of the manuscript. We thank Jonathan Lenz and Igor Mačinković for Schneider S2 cells. Funding was provided by Philipps University Marburg, Department of Medicine (to H.-R.C.).

## Author contributions

L.Z., J.G., H.S., C.B., B.W., and T.A. planned and performed experiments, contributed reagents, and analyzed data. S.-A.F. and O.S. planned experiments. S.-A.F. optimised MST assays and analyzed data. S.A. performed gel filtration. A.N. and T.S. performed high-throughput sequencing. H.-R.C. planned experiments and analyzed data. T.A. drafted the manuscript. T.A. and H.-R.C. wrote the final manuscript.

## Disclosure of Conflicts of Interest

None.

## Materials & correspondence

The corresponding authors are T. Adhikary and H.-R. Chung. Please address material requests to T. Adhikary.

## Notes

https://www.ncbi.nlm.nih.gov/geo/query/acc.cgi?acc=GSE215916

### Competing Interest Statement

The authors have declared no competing interest.

### Summary of Updates

Minor clarifications and additional references.

## References

1. J. Condeelis and J. W. Pollard. Macrophages: obligate partners for tumor cell migration, invasion, and metastasis. Cell, 124:263–6, 2006. doi:10.1016/j.cell.2006.01.007.

2. F. Finkernagel, S. Reinartz, S. Lieber, T. Adhikary, A. Wortmann, N. Hoffmann, T. Bieringer, A. Nist, T. Stiewe, J. M. Jansen, U. Wagner, S. Müller-Brüsselbach, and R. Müller. The transcriptional signature of human ovarian carcinoma macrophages is associated with extracellular matrix reorganization. Oncotarget, 7(46):75339–52, 2016. doi:10.18632/oncotarget.12180.

3. T. Adhikary, A. Wortmann, F. Finkernagel, S. Lieber, A. Nist, T. Stiewe, U. Wagner, S. Müller-Brüsselbach, and R. Müller. Interferon signaling in ascites-associated macrophages is linked to a favorable clinical outcome in a subgroup of ovarian carcinoma patients. BMC Genomics, 18(1):243, 2017. doi:10.1186/s12864-017-3630-9.

4. Adelaja, B. Taylor, K. M. Sheu, Y. Liu, S. Luecke, and A. Hoffmann. Six distinct NFκB signaling codons convey discrete information to distinguish stimuli and enable appropriate macrophage responses. Immunity, 54(5):916–30, 2021. doi:10.1016/j.immuni.2021.04.011.

5. Hoffmann and D. Baltimore. Circuitry of nuclear factor κB signaling. Immunological Reviews, 210:171–86, 2006. doi:10.1111/j.0105-2896.2006.00375.x.

6. S. Reinartz, F. Finkernagel, T. Adhikary, V. Rohnalter, T. Schumann, Y. Schober, A. W. Nockher, A. Nist, T. Stiewe, J. M. Jansen, U. Wagner, S. Müller-Brüsselbach, and R. Müller. A transcriptome-based global map of signaling pathways in the ovarian cancer microenvironment associated with clinical outcome. Genome Biol, 17:108, 2016. doi:10.1186/s13059-016-0956-6.

7. W. Dobryszycka. Biological functions of haptoglobin. Eur J Clin Chem Clin Biochem, 35(9):647–54, 1997.

8. M. Kristiansen, J. H. Graversen, C. Jacobsen, O. Sonne, H.-J. Hoffman, S. K. A. Law, and S. K. Moestrup. Identification of the haemoglobin scavenger receptor. Nature, 409(6817):198–201, 2001. doi:10.1038/35051594.

9. M. S. Arredouani, A. Kasran, J. A. Vanoirbeek, F. G. Berger, H. Baumann, and J. L. Ceuppens. Haptoglobin dampens endotoxin-induced inflammatory effects both in vitro and in vivo. Immunology, 114(2):263–71, 2005. doi:10.1111/j.1365-2567.2004.02071.x.

10. J. D. Belcher, C. Chen, J. Nguyen, F. Abdulla, P. Zhang, H. Nguyen, P. Nguyen, T. Killeen, S. M. Miescher, N. Brinkman, K. A. Nath, C. J. Steer, and G. M. Vercellotti. Haptoglobin and hemopexin inhibit vaso-occlusion and inflammation in murine sickle cell disease: Role of heme oxygenase-1 induction. PLoS One, 13(224):e0196455, 2018. doi:10.1371/journal.pone.0196455.

11. G. Galicia, W. Maes, G. Verbinnen, A. Kasran, D. Bullens, M. Arredouani, and J. L. Ceuppens. Haptoglobin deficiency facilitates the development of autoimmune inflammation. Eur J Immunol, 39(12):3404–12, 2009. doi:10.1002/eji.200939291.

12. H. Shen, Y. Song, C. M. Colangelo, T. Wu, C. Bruce, G. Scabia, A. Galan, M. Maffei, and D. R. Goldstein. Haptoglobin activates innate immunity to enhance acute transplant rejection in mice. J Clin Invest, 122(1), 2011. doi:10.1172/JCI58344.

13. J.-O. Kwon, W. J. Jin, B. Kim, H. Ha, H.-H. Kim, and Z. H. Lee. Haptoglobin Acts as a TLR4 Ligand to Suppress Osteoclastogenesis via the TLR4–IFN-β Axis. J Immunol, 202(12):3359–69, 2019. doi:10.4049/jimmunol.1800661.

14. P. Boroni Moreira, T. F. Salles Texeira, A. Barbosa Ferreira, M. do Carmo Gouveia Peluzio, and R. de Cássia Gonçalves Alfenas. Influence of a high-fat diet on gut microbiota, intestinal permeability and metabolic endotoxaemia. Br J Nutr, 108(5):801–9, 2012. doi:10.1017/S0007114512001213.

15. K. de Punder and L. Pruimboom. Stress induces endotoxemia and low-grade inflammation by increasing barrier permeability. Front Immunol, 6:223, 2015. doi:10.3389/fimmu.2015.00223.

16. C. Erridge, T. Attina, C. M. Spickett, and D. J. Webb. A high-fat meal induces low-grade endotoxemia: evidence of a novel mechanism of postprandial inflammation. Am J Clin Nutr, 86(5):1286–92, 2007. doi:10.1093/ajcn/86.5.1286.

17. M. I. Lassenius, K. H. Pietiläinen, K. Kaartinen, P. J. Pussinen, J. Syrjänen, C. Forsblom, I. Pörsti, A. Rissanen, J. Kaprio, J. Mustonen, P.-H. Groop, M. Lehto, and FinnDiane Study Group. Bacterial Endotoxin Activity in Human Serum Is Associated With Dyslipidemia, Insulin Resistance, Obesity, and Chronic Inflammation. Diabetes Care, 34(8):1809–15, 2011. doi:10.2337/dc10-2197.

18. H. Fukui, B. Brauner, J. C. Bode, and C. Bode. Plasma endotoxin concentrations in patients with alcoholic and non-alcoholic liver disease: reevaluation with an improved chromogenic assay. J Hepatol, 12(2):162–9, 1991. doi:10.1016/0168-8278(91)90933-3.

19. M. D. Nguyen, J.-P. Julien, and S. Rivest. Innate immunity: the missing link in neuroprotection and neurodegeneration? Nat Rev Neurosci, 3(3):216–27, 2002. doi:10.1038/nrn752.

20. G. C. Brown. The endotoxin hypothesis of neurodegeneration. JNI, 16(1):180, 2019. doi:10.1186/s12974-019-1564-7.

21. R. F. Schwabe and C. Jobin. The microbiome and cancer. Nat Rev Cancer, 13(11):800–12, 2013. doi:https://doi.org/10.1038/nrc3610.

22. J. M. Moreno-Navarrete and J. M. Fernández-Real. Antimicrobial-Sensing Proteins in Obesity and Type 2 Diabetes. Diabetes Care, 34(Suppl 2):S335–41, 2011. doi:10.2337/dc11-s238.

23. M. M. Wurfel, B. G. Monks, R. R. Ingalls, R. L. Dedrick, R. Delude, D. Zhou, N. Lamping, R. R. Schumann, R. Thieringer, M. J. Fenton, S. D. Wright, and D. Golenbock. Targeted deletion of the lipopolysaccharide (LPS)-binding protein gene leads to profound suppression of LPS responses ex vivo, whereas in vivo responses remain intact. J Exp Med, 186(12):5051–6, 1997. doi:10.1084/jem.186.12.2051.

24. J.C. Marshall. Lipopolysaccharide: An Endotoxin or an Exogenous Hormone? Clinical Infectious Diseases, 41 (Suppl. 7):S470–80, 2005. doi:10.1086/432000.

25. K. A. Fitzgerald, D. C. Rowe, B. J. Barnes, D. R. Caffrey, A. Visintin, E. Latz, B. Monks, P. M. Pitha, and D. T. Golenbock. LPS-TLR4 Signaling to IRF-3/7 and NF-ŒR BInvolvestheTollAdaptersTRAMandTRIF. J Exp Med, 198 (7) : 1043–-55, 2003. doi : 10.1084/jem.20031023.

26. C.-M. Tsai and C. E. Frasch. A Sensitive Silver Stain for Detecting Lipopolysaccharides in Polyacrylamide Gels. Anal Biochem, 119(1):115–9, 1981. doi:10.1016/0003-2697(82)90673-x.

27. A. P. Morgan, I. M. Helander, and T. U. Kosunen. Compositional analysis of Helicobacter pylori Rough-Form Lipopolysaccharides. J Bacteriol, 174(4):1370–7, 1992. doi:10.1128/jb.174.4.1370-1377.1992.

28. S. C. Bailey and D. Apirion. Identification of Lipopolysaccharides and Phospholipids of Escherichia coli in Polyacrylamide Gels. J Bacteriol, 131(1):347–55, 1977.

29. W. Kaca, R. I. Roth, and J. Levin. Hemoglobin, a newly recognized lipopolysaccharide (LPS)-binding protein that enhances LPS biological activity. J Biol Chem, 269(40):25078–84, 1994. doi:10.1016/S0021-9258(17)31501-6.

30. T. Ogawa, Y. Asai, Y. Makimura, and R. Tamai. Chemical structure and immunobiological activity of Porphyromonas gingivalis lipid A. Front Biosci (Landmark Ed), 12(10):3795–812, 2007. doi:10.2741/2353.

31. V. Bagaev, A. Y. Garaeva, E. S. Lebedeva, A. V. Pichugin, R. I. Ataullakhanov, and F. I. Ataullakhanov. Elevated pre-activation basal level of nuclear NF-κB in native macrophages accelerates LPS-induced translocation of cytosolic NF-κB into the cell nucleus. Sci Rep, 9(1):4563, 2019. doi:10.1038/s41598-018-36052-5.

32. Q. J. Cheng, S. Ohta, K. M. Sheu, R. Spreafico, A. Adelaja, B. Taylor, and A. Hoffmann. NF-kB dynamics determine the stimulus specificity of epigenomic reprogramming in macrophages. Science, 372(6548):1349–53, 2021. doi:10.1126/science.abc0269.

33. M. Son, A. G. Wang, H.-L. Tu, M. Oliver Metzig, P. Patel, K. Husain, J. Lin, A. Murugan, A. Hoffmann, and S. Tay. NF-κB responds to absolute differences in cytokine concentrations. Science Signaling, 14(666):eaaz4382, 2021. doi:10.1126/scisignal.aaz4382.

34. J. Schaer, P. W. Buehler, A. I. Alayash, J. D. Belcher, and G. M. Vercellotti. Hemolysis and free hemoglobin revisited: exploring hemoglobin and hemin scavengers as a novel class of therapeutic proteins. Blood, 121(8), 2013. doi:10.1182/blood-2012-11-451229.

35. G. Petruk, M. Puthia, J. Petrlova, F. Samsudin, A.-C. Strömdahl, S. Cerps, L. Uller, S. Kjellström, P. J. Bond, and A. Schmidtchen. SARS-CoV-2 spike protein binds to bacterial lipopolysaccharide and boosts proinflammatory activity. J Mol Cell Biol, 12(12):916–32, 2020. doi:10.1093/jmcb/mjaa067.

36. S. Tumpara, A. R. Gründing, K. Sivaraman, S. Wrenger, B. Olejnicka, T. Welte, M. J. Wurm, P. Pino Kiseljak, F. M. Wurm, and S. Janciauskiene. Boosted Pro-Inflammatory Activity in Human PBMCs by Lipopolysaccharide and SARS-CoV-2 Spike Protein Is Regulated by α-1 Antitrypsin. Int J Mol Sci, 22(15):7941, 2021. doi:10.3390/ijms22157941.

37. K. Kruglikov and P. E. Scherer. Preexisting and inducible endotoxemia as crucial contributors to the severity of COVID-19 outcomes. PLOS Pathogens, 17(2):e1009306, 2021. doi:10.1371/journal.ppat.1009306.

38. S. F. Assimakopoulos, S. Mastronikolis, A.-L. de Lastic, D. Aretha, D. Papageorgiou, T. Chalikidi, I. Oikonomou, C. Triantos, A. Mouzaki, and M. Marangos. Intestinal Barrier Biomarker ZO1 and Endotoxin Are Increased in Blood of Patients With COVID-19-associated Pneumonia. In Vivo, 35(4), 2021. doi:10.21873/invivo.12528.

39. A. Tripathi, K. M. Lammers, S. Goldblum, T. Shea-Donahue, S. Netzel-Arnett, M. S. Buzza, T. M. Antalis, S. N. Vogel, A. Zhao, S. Yang, M.-C. Arrietta, J. B. Meddings, and A. Fasano. Identification of human zonulin, a physiological modulator of tight junctions, as prehaptoglobin-2. Proc Natl Acad Sci, 106(39):39, 2009. doi:10.1073/pnas.0906773106.

40. D. Su, R. I. Roth, M. Yoshida, and J. Levin. Hemoglobin increases mortality from bacterial endotoxin. Infect Immun, 65(4):1258–66, 1997. doi:10.1128/iai.65.4.1258-1266.1997.

41. T. Lin, Y. H. Kwak, F. Sammy, P. He, S. Thundivalappil, G. Sun, W. Chao, and H. S. Warren. Synergistic inflammation is induced by blood degradation products with microbial Toll-like receptor agonists and is blocked by hemopexin. J Infect Dis, 202(4):624–32, 2010. doi:10.1086/654929.

42. M. Nakamura, H. J. Bax, D. Scotto, E. Amiri Souri, S. Sollie, R. J. Harris, N. Hammar, G. Walldius, A. Winship, S. Ghosh, A. Montes, J. F. Spicer, M. van Hemelrijck, D. H. Josephs, K. E. Lacy, S. Tsoka, and S. N. Karagiannis. Immune mediator expression signatures are associated with improved outcome in ovarian carcinoma. OncoImmunology,8(6), 2019. doi:10.1080/2162402X.2019.1593811.

43. C.-S. Tai, Y.-R. Lin, T.-H. Teng, P.-Y. Lin, S.-J. Tu, Y.-R. Huang, W.-C. Huang, S.-L. Weng, H.-D. Huang, Y.-L. Chen, and W. L. Chen. Haptoglobin expression correlates with tumor differentiation and five-year overall survival rate in hepatocellular carcinoma. PLoS One, 12(2):e0171269, 2017. doi:10.1371/journal.pone.0171269.

44. S.-K. Oh, D. L. Very, J. Walker, S Raam, and S. T. Ju. An analogy between fetal haptoglobin and a potent immunosuppressant in cancer. Cancer Res, 47:5120–6, 1987.

45. N. Okuyama, Y. Ide, M. Nakano, T. Nakagawa, K. Yamanaka, K. Morikawi, K. Murata, H. Ohigashi, S. Yokoyama, H. Eguchi, O. Ishikawa, T. Ito, M. Kato, A. Kasahara, S. Kawano, J. Gu, N. Taniguchi, and E. Miyoshi. Fucosylated haptoglobin is a novel marker for pancreatic cancer: A detailed analysis of the oligosaccharide structure and a possible mechanism for fucosylation. Int J Cancer, 118(11):2803–8, 2006. doi:10.1002/ijc.21728.

46. A. Unger, F. Finkernagel, N. Hoffmann, F. Neuhaus, B. Joos, A. Nist, T. Stiewe, A. Visekruna, U. Wagner, S. Reinartz, S. Müller-Brüsselbach, R. Müller, and T. Adhikary. Chromatin binding of c-REL and p65 is not limiting for macrophage IL12B transcription during immediate suppression by ovarian carcinoma ascites. Front Immunol, 9:1425, 2018. doi:10.3389/fimmu.2018.01425.

47. S.-A. Freibert, M. T. Boniecki, C. Stümpfig, V. Schulz, N. Krapoth, D. R. Winge, U. Mühlenhoff, O. Stehling, M. Cygler, and R. Lill. N-terminal tyrosine of ISCU2 triggers [2Fe-2S] cluster synthesis by ISCU2 dimerization. Nat Commun, 12(1), 2021. doi:10.1038/s41467-021-27122-w.

